# *In Silico* Identification, Characterization and Diversity Analysis of RNAi Genes and their Associated Regulatory Elements in Sweet Orange (*Citrus sinensis L*)

**DOI:** 10.1101/2020.01.13.904128

**Authors:** Md. Parvez Mosharaf, Md. Asif Ahsan, Hafizur Rahman, Zobaer Akond, Fee Faysal Ahmed, Md. Mazharul Islam, Mohammad Ali Moni, Md. Nurul Haque Mollah

## Abstract

RNA interference (RNAi) plays key roles in post-transcriptional and chromatin modification levels as well as regulates various eukaryotic gene expressions which involved in stress responses, development and maintenance of genome integrity during developmental stages. The whole mechanism of RNAi pathway is directly involved with the gene-silencing process by the interaction of Dicer-Like (DCL), Argonaute (AGO) and RNA-dependent RNA polymerase (RDR) gene families. However, the genes of these three RNAi families are largely unknown yet in sweet orange (*Citrus sinensis)*, though it is an economically important fruit plant all over the world. Therefore, a comprehensive investigation for genome-wide identification, characterization and diversity analysis of RNA silencing genes in *C. sinensis* was conducted and identified 4 CsDCL, 8 CsAGO and 4 CsRDR as RNAi genes. To characterize and validate the predicted genes of RNAi families, various bioinformatics analysis was conducted. Phylogenetic analysis clustered the predicted CsDCLs, CsAGOs and CsRDRs genes into four, six and four subgroups with the relevant genes of Arabidopsis respectively. The domain and motif composition analysis, the gene structure for all three-gene families exhibited almost homogeneity within the same group members while showed significant differences in between groups. The GO enrichment analysis results clearly indicated that the predicted genes have direct involvement into the RNAi process as expected in *C. sinensis*. Moreover, *Cis*-regulatory elements and regulatory transcription factor analysis of the reported RNAi genes demonstrated the diverse connection to the huge biological functions and regulatory pathways. The expressed sequence tag (EST) analysis showed that these genes are highly expressed in fruit and leaves which indicate that these reported genes have great involvement in *C. sinensis* food, flowering and fruit production. The expression analysis of the reported RNAi genes might be more useful to explore the most effective RNAi genes in *C. sinensis* for further biotechnological application.

## 1. INTRODUCTION

In multicellular eukaryotes, wide range of biological functions including genome rearrangement, antiviral defense, heterochromatin formation and development patterning and timing are fine-tuned by generally two types of small RNA (sRNA; including 21-24 nucleotides), named microRNA (miRNA) and short interfering RNA (siRNA) [1–3]. These sRNA molecules are initially involved with both transcriptional silencing and post-transcriptional RNA interfering RNA formation [2]. In plants, the sRNA biogenesis process is significantly regulated by the proteins encoded by respective Dicer-like (DCL), Argonate (AGO) and RNA-dependent RNA polymerases (RDR) gene families. The DCLs are deeply related to process the complementary dsRNAs into sRNAs especially 21-24 nucleotides longer (i.e. siRNA or miRNA). The specification to the endonuclease-containing, RNA-induced silencing complex (RISC) is provided by these sRNAs which facilitate the AGO proteins with RNaseH-type activities to degrade the target homologous RNAs with the sequence complementary to the small RNAs. These also refer to the transcriptional gene silencing by the implementation of chromatin reformation [4, 5].

A major component of RNA interference (RNAi) pathway (also known as small RNA (sRNA) biogenesis process), called DCL proteins which mainly process the small mature RNAs from the long double-stranded RNAs [6–10]. The important characteristic of these DCL proteins that having the functional domains, named DEAD/ResIII, Helicase_C, Dicer_Dimer, PAZ, RNase III and DSRM [11]. The PAZ domain acts to bind the siRNA as well as the dsRNA is cleaved by the two catalytic RNaseIII domains. The main components of RNAi are the AGO proteins which play the core roles of RNA silencing [12]. All the AGO proteins include the Argo-N/Argo-L, PAZ, MID and PIWI significant functional domains [13]. A significant specific binding pocket is contained in the PAZ domain. Additionally, to anchor the sRNA onto the AGO proteins, the specific pocket of MID domain binds the 5′ phosphate of the small RNAs [14]. The siRNA5′ end is bonded to the target RNA by the PIWI domain [15]. Among the three groups of AGO proteins i.e. Ago -like, PIWI-like and *C. elegens*-specific group 3 AGO proteins [16, 17], the Ago-like proteins are presented and expressed in plants, animals, fungi and bacteria, while PIWI-like proteins have only found in animals [18]. Some important catalytic residues are missed by *C. elegens-*specific group 3 AGO proteins [16] but others conserved as well as the expression of PIWI-like group is restricted in human germ-cell and in rat and some mammals [18]. The third major RNAi protein is RDR which have not been identified in insects or vertebrates [19] but present in fungi, nematodes and plants. These proteins are closely related to the various types of gene silencing [13].

In case of plants, multiple genes belong to the DCL, AGO and RDR gene families related to distinct RNAi pathways [20–22]. The member of these RNAi gene families has been identified in many plants species (such as Arabidopsis [23], rice [13], maize [24], tomato [25], tobacco [26], foxtail millet [27], and grapevine [28], cucumber [29], pepper [30]). In Arabidopsis, AtDCL1, AtDCL3 and AtAGO4 influenced the RNA-directed DNA methylation of the *FWA* transgene linkage to the histone H3 lysine 9 (H3K9) methylation [31, 32]. AtDCL2 is associated with the virus defence and siRNA production when the AtDCL4 is related to the regulation of vegetative phase change [11, 33]. AtDCL1 and AtDCL3 function for Arabidopsis flowering [34]. Moreover, in rice if the OsDCL1 is knocked down then it fails to perform siRNA metabolism which results pleiotropic phenotype in rice [35].

Besides, AGO proteins related to various forms of RNA silencing, such as AtAGO1 is associated with the transgene-silencing pathways [36] and AtAGO4 with the epigenetic silencing [31]. AtAGO7 and AtAGO10 influence the plant growth [37] and meristem maintenance [38]. Additionally, others AGOs also have a significant role in RNAi pathways. On the other hand, previous studies reported that the RDR genes are responsible for different gene silencing including co-suppression, virus defence, chromatin silencing and PTGS in plants such as in Arabidopsis, maize [21, 39–42].

However, the genes of these three RNAi families are largely unknown yet in the economically important fruit plant *C. sinensis.* The sweet orange (*C. sinensis*) is considered as an inevitable source of high nutrition, a great natural source of vitamin C, an unavoidable antioxidant important for the human body [43]). In production volume, orange is the world second fruit plant all over the world (FAO Statistics 2006). Not only having the market value, but also sweet orange contains about 170 phytonutrients and over 60 flavonoids those works as antioxidant, anti-inflammation, anti-cancer and anti-arteriosclerosis compound and also protects from many chronic diseases like arthritis, obesity and coronary heart diseases [44–47]. Therefore, a comprehensive investigation for genome-wide identification, characterization and diversity analysis of RNAi genes in *C. sinensis* is essential. In this project, an attempt is made to accomplish a comprehensive *in silico* investigation for genome-wide identification, characterization and diversity analysis of the members of DCL, AGO and RDR gene families in *C. sinensis*. A number of bioinformatics analysis such as, phylogenetic analysis, gene structure, functional domains and motifs, gene ontology (GO), sub-cellular localization, transcription factor (TF) analysis, cis-acting regulatory element and expressed sequence tag (EST) analysis have been considered to characterize and validate the predicted RNAi genes in *C. sinensis*.

## 2. MATERIALS AND METHODS

### 2.1. Identification of DCL, AGO and RDR Genes

For genome-wide identification of DCL, AGO and RDR genes in sweet orange (*C. sinensis*), protein sequences were downloaded from the Phytozome database (https://phytozome.jgi.doe.gov/pz/portal.html). In this purpose, the previously identified sRNA biogenesis protein sequences of the model plant *Arabidopsis thaliana* (AtDCLs, AtAGOs and AtRDRs) were used to search the protein sequence of *C. sinensis*. The Basic Local Alignment Search Tool (BLASTP) program was used against *C. sinensis* genome in the Phytozome database (Fig. **(1)**).

**Fig.(1).**
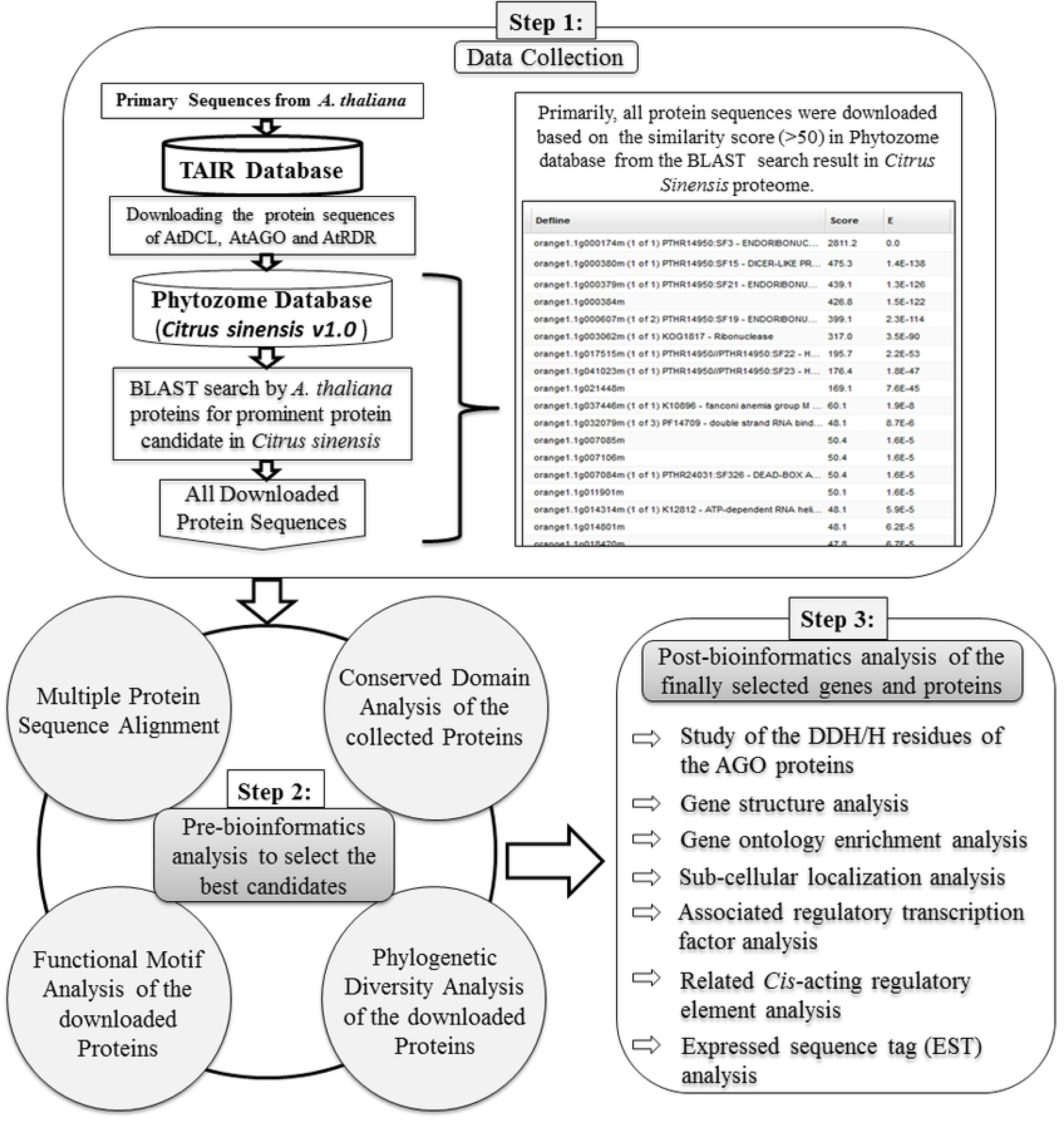
The working flowchart to select the best candidates for DCL, AGO and RDR genes in *C. sinensis*.

The resulted paralogs protein sequences from *C. sinensis* were downloaded with the significant score (≥50) and the significant E-values. For avoiding the redundancy of sequences, only the primary transcripts were considered in this analysis. The conserved domains of all retrieved sequences were searched and predicted by using the Pfam (http://pfam.sanger.ac.uk/) and the NCBI-CD database (http://www.ncbi.nlm.nih.gov/Structure/cdd/wrpsb.cgi) and the SMART analysis. By this time, the different genomic information including the primary transcript name, genomic length, the chromosomal location of genes, encoded protein length were downloaded from the *C*. *sinensis* genome in Phytozome database. In this study, the computationally identified new CsDCLs, CsAGOs and CsRDRs genes in *C. sinensis* genome were named according to the nomenclature based on phylogenetic relatedness of the similar family-members of the *Arabidopsis thaliana* genes as named previously. The molecular weight of the selected protein sequences was predicted by using the ExPASyComputepI/Mwtool (http://au.expasy.org/tools/pitool.html).

### 2.2. Sequence Alignment and Phylogenetic Analysis

In this *in silico* identification, the multiple sequence alignments of the encoded protein sequences of the predicted genes were conducted by using the Clustal-W method [48] with the MEGA5 program [49]. Finally, the phylogenetic tree analysis was carried out using the Neighbor-joining method [50] implemented on the aligned sequenced and the 1,000 bootstrap replicates [51] were used to check this evolutionary relationship. The evolutionary distances were computed using the Equal Input method [52].

### 2.3. Conserved Domain and Motif Analysis

To investigate the functional domains of the predicted genes by GENEDOC program using the aligned sequences although domains were predicted according to the NCBI database, Pfam database and SMART analysis (Fig. **(1)**). The CsDCLs, CsAGOs and CsRDRs proteins reflected multiple functional domains like the AtDCLs, AtAGOs and AtRDRs proteins. The predicted genes were selected containing the maximum number of functional domains similar to the AtDCLs, AtAGOs and AtRDRs. For DCL proteins, all six functional domains i.e. DEAD/ResIII, Helicase_C, Dicer_Dimer, PAZ, RNase III and DSRM were considered to choose the prominent CsDCL candidates in *C*. *sinensis.* Similarly, the proteins were sorted out as CsAGO candidate having the Argo-N/Argo-L, PAZ, MID and PIWI significant functional domains, the characteristic of plant AGO proteins. Same approach was applied to discriminate the CsRDRs genes. In motif investigation, the most significant conserved metal-chelating catalytic triad residues in the PIWI domain, i.e. aspartate, aspartate and histidine (DDH) [13] as well as histidine at 798 positions (H798) were considered for CsAGOs (Fig. **(2)**). In this study, we considered the alignment of CsAGOs with the paralogs AtAGOs using Clustal-W method as mentioned before. Furthermore, the conserved motif divergences among all the predicted genes were conducted by a complete online protein sequence analysis using Multiple Expectation Maximization for Motif Elicitation (MEME-Suite) [53]. For this purpose, the following parameters were specified: (i) optimum motif width as ≥6 and ≤50; (ii) maximum 20 motifs.

**Fig. (2).**
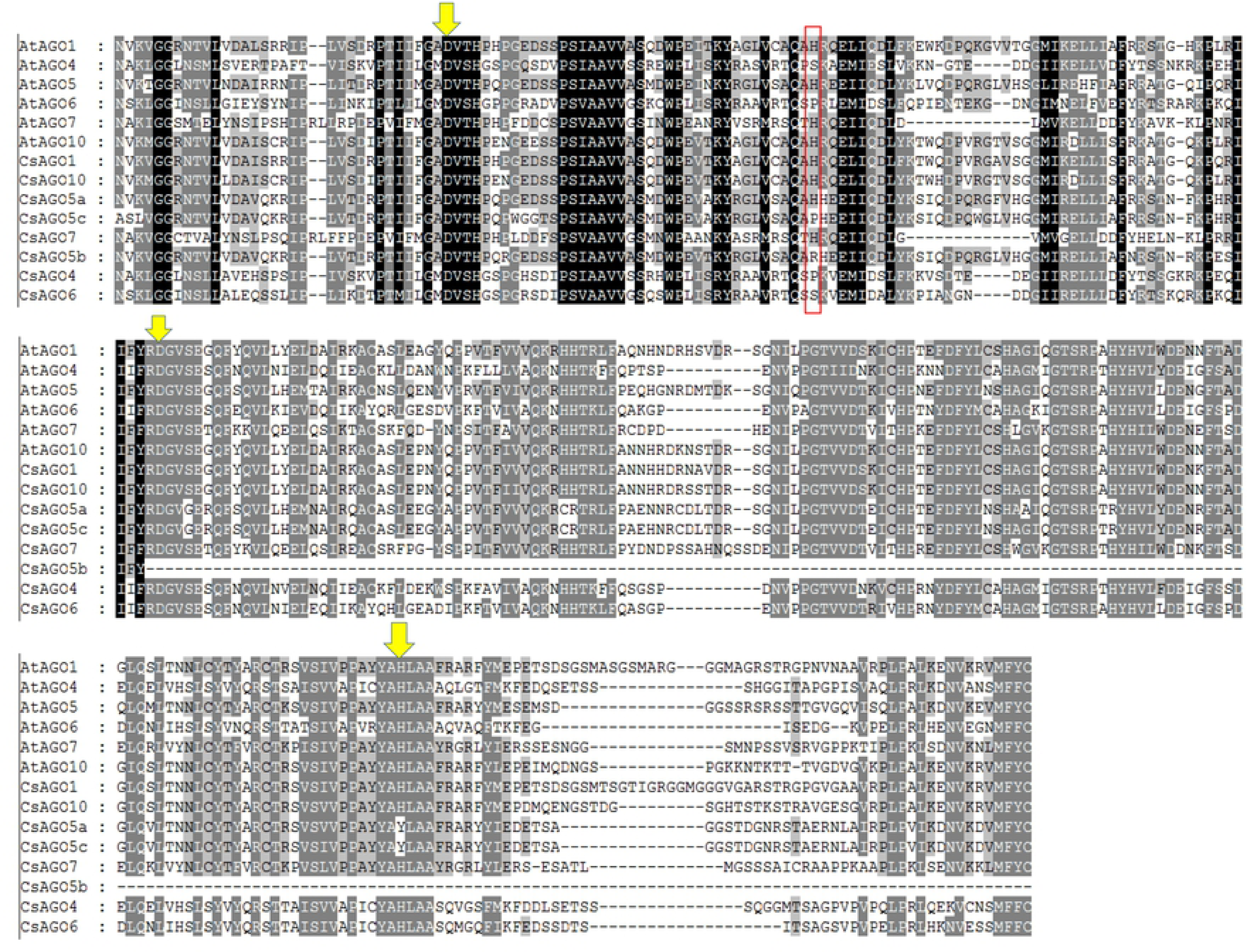
The multiple sequence alignment profile of PIWI domain of the amino acids sequences of *C. sinensis* and Arabidopsis AGO proteins by Clustal-W program in MEGA5. The downward yellow arrows indicate the position of conserved DDH traid of PIWI domain and the conserved H798 positions are surrounded by red box.

### 2.4. Gene Structure Analysis

For further computational analysis, we have considered the predicted gene structure and their interaction network analysis (Fig. **(1)**). The gene structure was analyzed using the online Gene Structure Display Server (GSDS 2.0, http://gsds.cbi.pku.edu.cn/index.php). The structures of the selected genes were compared with the gene structure of *Arabidopsis thaliana.* These combined gene structure Figures revealed the exon-intron composition of the predicted gene in *C*. *sinensis.* Moreover, numbers of intron were noted from GSDS.

### 2.5. Gene Ontology and Sub-cellular Localization Analysis

To check the engagement of our predicted RNAi associated genes with the cluster of different biological processes and molecular functional pathways, the GO analysis was conducted using online tool implemented in PlantTFDB[54]. Here, the respective p-values were determined by Fisher’s exact test and Benjamini-Hochberg’s technique was chosen for multiple testing corrections. We considered the p-value < 0.05 as statistical significant for the GO enrichment results corresponding to the predicted genes. For the reported gene product, the sub-cellular location was investigated into the cell considering the different organelles (Fig. **(1)**). these allocations could improve the overall understanding of the proposed genes as well as proteins functional pathways through the RNAi process. Web-based integrative subcellular location predictor tool called plant subcellular localization integrative predictor (PSI) [55] was used to predict the subcellular location of the identified genes.

### 2.6. Identification of Regulatory Relationship between TFs and *C. sinensis* RNAi genes

In every living organism, transcription levels of genes are directly regulated by the expression or suppression activities of transcription factors (TF) associated with the respective genes. TFs are also a kind of protein which can bind to the specific promoter region of the genes and can control the transcription process whether the genes to be expressed or suppressed. TFs are involved in the plant stress response activities; plant development in different life stage, disease resistance and so on. In this study, the analysis of associated TFs family with the predicted RNAi genes in *C*. *sinensis* was conducted from the widely used plant transcription factor database, plantTFDB (http://planttfdb.cbi.pku.edu.cn//). The related TFs were also categorized into 27 different TFs families and their roles in plants are also provided. These TFs are extremely important to study for the development of *C*. *sinensis* plant and enrich the sweet orange production.

### 2.7. Construction of Regulatory Network between TFs and *C. sinensis* RNAi genes

After identification of the related regulatory TFs of the *C. sinensis* RNAi genes, the regulatory network and sub-network were constructed and visualized using Cytoscape 3.7.1 [56] to find out the hub proteins and the related important TF through the interaction network. The networks were constructed to investigate the regulatory relationship between the TFs and RNAi genes.

### 2.8. *Cis*-regulatory element Analysis

In genetic transcription, the *cis*-elements are highly associated by controlling the transcription of a specific gene through the regulatory transcription factors. To collect more related genomic information about the *C. sinensis* RNAi genes, the CIs-acting regulatory element In the promoter region was retrieved and analyzed through the online c*is*-element prediction database PlantCARE (http://bioinformatics.psb.ugent.be/webtools/plantcare/html/). The collected *cis-*regulatory element was categorized into five categories like light responsive (LR), stress responsive (SR), hormone responsive (HR), others activities (OT) and unknown function. The cis-element having known and described function are represented into the Fig. 12 for CsDCLs, CsAGOs and CsRDR separately.

### 2.9. *In silico* Expressed Sequence Tag (EST) Analysis

For the important and valuable information about the gene expression, the *in silico* expressed sequence tag (EST) data analysis was conducted. The PlantGDB (http://www.plantgdb.org/cgi-bin/blast/PlantGDB/) was used for EST mining against the proposed RNAi genes in *C. sinensis*. The default parameter with e-value=1e-10 were considered for blastn search for the EST mining in PlantGDB database. The further heatmap was constructed to represent the specific RNAi gene expression into different tissue and organ in this fruit plant.

## 3. RESULTS AND DISCUSSION

### 3.1. Identification and characterization of CsDCLs, CsAGOs and CsRDRs genes

To identify the best candidates of RNAi pathway in *C*. *sinensis* similar to the *A. thaliana*, all the previously downloaded sequences were gone throughout various kinds of analysis (Fig. **(1)**). Finally, we have isolated 4 DCL genes, 8 AGO genes and 4 RDR genes those produce the CsDCLs proteins, CsAGOs and CsRDRs proteins respectfully in the *C. sinensis* genome. On the basis of HMMER analysis with regards of all six types of conserved domains DEAD, Helicase_C, Dicer_dimar, PAZ, RNase III and DSRM; four DCL loci were identified in sweet orange genome. From the conserved domain search by Pfam databases, NCBI databases and the SMART analysis, all reflected that half of the predicted proteins (CsDCL1 and CsDCL4) are conserved with the DEAD/ResIII, Helicase_C, Dicer-dimer, PAZ, RNase III and dsRM domains, which are preserved by all the plant DCL proteins from the DCL genes family (class 3 RNase III family) [57, 58]. On the other hand, CsDCL2 and CsDCL3 have missed a single DSRM domain while others are contained with a second DSRM domain. This DsRM domain is lacked in by non-plant DCL completely [11]. Compared to others DCL proteins, the CsDCL1 has the N-terminal DEAD domain which might consist of three adjacent segments of amino acid sequence within the full domain length (152 amino acids), resulted by the analysis of Pfam databases and SMART. The CsDCL3 also revealed the ResIII domain instead of the DEAD domain. Additionally, the genome length of predicted CsDCL genes varied from 10603 bp to 12728 bp corresponding to CsDCL1 and CsDCL2 with the coding potentiality of 1931 and 1396 amino acid (Table **(1)**).

**Table 1:** Basic information about *C*. *sinensis* Dicer-like, Argonaute and RNA-dependent RNA polymerase gene families.

| Serial Number | Gene Name | Accession Number | Chromosomal location | Gene Length (bp) | Protein |  |  |  | Type |
| --- | --- | --- | --- | --- | --- | --- | --- | --- | --- |
|  |  |  |  |  | No. of Intron | Molecular Weight (D.a) | Protein Length (a.a) | PI |  |
| CsDCLs |  |  |  |  |  |  |  |  |  |
| 1 | CsDCL1 | orange1.lg000174m | scaffold00001:3480331..3490933 | 10603 | 18 | 216424.20 | 1931 | 5.96 | DCL 1 |
| 2 | CsDCL2 | orange1.lg000607m | scaffold00367:74912..87639 | 12728 | 21 | 158487.91 | 1396 | 7.65 | DCL 2 |
| 3 | CsDCL3 | orange1.lg000379m | scaffold00013:998934..1009643 | 10710 | 24 | 178949.30 | 1604 | 8.01 | DCL 3 |
| 4 | CsDCL4 | orange1.lg000380m | scaffold00068:219006..230745 | 11740 | 25 | 179592.47 | 1601 | 6.46 | DCL 4 |
| CsAGOs |  |  |  |  |  |  |  |  |  |
| 1 | CsAGO1 | orange1.lg001466m | scaffold00674:51962..59926 | 7965 | 20 | 118338.09 | 1073 | 9.38 | AGO 1 |
| 2 | CsAGO4 | orange1.lg002449m | scaffold00028:907375..916121 | 8747 | 21 | 102997.53 | 920 | 8.98 | AGO 4 |
| 3 | CsAGO5a | orange1.lg002204m | scaffold00595:85931..92382 | 6452 | 21 | 106772.65 | 954 | 9.27 | AGO 5 |
| 4 | CsAGO5b | orange1.lg037086m | scaffold00595:96826..99593 | 2768 | 10 | 48003.28 | 426 | 9.05 | AGO 5 |
| 5 | CsAGO5c | orange1.lg003630m | scaffold03700:12..6517 | 6506 | 21 | 91163.31 | 806 | 9.03 | AGO 5 |
| 6 | CsAGO6 | orange1.lg002661m | scaffold00067:338566..346634 | 8069 | 21 | 100499.54 | 895 | 9.41 | AGO 6 |
| 7 | CsAGO7 | orange1.lg001684m | scaffold00003:3309646..3313553 | 3908 | 2 | 117035.78 | 1030 | 9.24 | AGO 7 |
| 8 | CsAGO10 | orange1.lg001954m | scaffold00011:54251..63917 | 9667 | 20 | 111515.83 | 992 | 9.34 | AGO 10 |
| CsRDRs |  |  |  |  |  |  |  |  |  |
| 1 | CsRDR1 | orange1.lg002586m | scaffold00058:231382..236081 | 4700 | 2 | 103382.03 | 904 | 6.51 | RDR 1 |
| 2 | CsRDR2 | orange1.lg001183m | scaffold00020:1255446..1250927 | 4520 | 3 | 128958.15 | 1131 | 6.59 | RDR 2 |
| 3 | CsRDR3 | orange1.lg001771m | scaffold00027:638984..650509 | 11526 | 18 | 115670.83 | 1015 | 6.80 | RDR 3 |
| 4 | CsRDR6 | orange1.lg041430m | Scaffold d00051:398116..402488 | 4373 | 1 | 136437.20 | 1157 | 6.13 | RDR 6 |

[Information containing the gene names, accession number, chromosomal location, genome length, protein length were collected from Phytozome database (https://phytozome.jgi.doe.gov/pz/portal.html) and the molecular weight and PI values were predicted by the ExPASy ComputePI/Mwtool (http://au.expasy.org/tools/pi_tool.html). Molecular weights are in Daltons and “aa” means amino acid.]

Based on the conserved domain PAZ and PIWI from the putative polypeptide sequences by HMM and HMMER analysis, we have isolated a total of 8 AGO genes in the *C. sinensis* genome. Conserved domain analysis by the Pfam database, NCBI databases and SMART analysis, reported that all the selected AGO proteins (CsAGO1-8) shared an N-terminus PAZ domain and a C-terminus PIWI super family domain, the core properties of plant AGO proteins. Moreover, the CsAGO proteins also preserved the other domains like the Arabidopsis i.e. ArgoN, ArgoL1, DUF1785, ArgoL2, ArgoMid.

From the previous study, the PIWI domain demonstrating expansive homology to RNase H binds the siRNA 5′ end to the target RNA [15] and cracks the target RNAs that represents the complementary sequences to small RNAs [59]. Interestingly, the catalytic traid, three conserved metal-chelating residues (D=aspartate, D=aspartate and H=histidine) in PIWI domain are related to the previous event and this traid was firstly showed in the model plant Arabidopsis on AGO1 [13].

Moreover, another critical conserved histidine residue in AGO1 for *in vitro* endonuclease activity [60] was found. The genome length of the selected CsAGO genes varied from 2768 bp to 9667 bp producing the CsAGO5b and CsAGO10 respectively with the coding potentiality of 426 and 992 amino acids. In this study, the multiple sequence alignment of the PIWI domains of all CsAGOs with the paralogs AtAGOs in Arabidopsis using the CLUSTAL-W method (Fig. **(2)**).

This alignment revealed that the five CsAGOs represented the conserved DDH traid residues like Arabidopsis AGO1. Among the other three CsAGOs, the CsAGO5b represented two missing PIWI domain catalytic residue(s) in the second aspartate at the 845^th^ position (D845) and third histidine at the 986^th^ position (H986) (Table **(2)**). But the other two CsAGO proteins, CsAGO5a and CsAGO5c has the catalytic traid but with a replacement in the third histidine residue at the 986^th^ position by tyrosine (Y) residue.

**Table 2:** Comparison of the argonaute proteins with missing catalytic residue(s) in PIWI domains between *C*. *sinensis* and *A. thaliana*.

| Serial No. | *Citrus sinensis* |  | *Arabidopsis thaliana*<sup>b</sup> |  |
| --- | --- | --- | --- | --- |
|  | Argonaute | Motifs<sup>a</sup> | Argonaute | Motifs<sup>a</sup> |
| 1 | CsAGO1 | DDH/H | AtAGO1 | DDH/H |
| 2 | CsAGO4 | DDH/P | AtAGO4 | DDH/S |
| 3 | CsAGO5a | DDY/H | AtAGO5 | DDH/H |
| 4 | CsAGO5b | D__/R | AtAGO6 | DDH/P |
| 5 | CsAGO5c | DDY/P | AtAGO7 | DDH/H |
| 6 | CsAGO6 | DDH/S | AtAGO10 | DDH/H |
| 7 | CsAGO7 | DDH/H |  |  |
| 8 | CsAGO10 | DDH/H |  |  |

<sup>a</sup>Comparison of conserved motif corresponds to D760, D845, H986/H798 of Arabidopsis AGO1; D: aspartate, H: histidine, P: proline, R: arginine, S: serine, Y: tyrosine and “\_” represents the missing catalytic residue(s).
<sup>b</sup>From Kapoor et al. (2008)

Surprisingly, the histidine residue at the 786^th^ position was replaced by proline (P) in CsAGO4 and CsAGO5c; by arginine (R) in CsAGO5b and in CsAGO6, H786 residue was replaced by serine (S) residue (Table **(2)**). The newly identified 4 CsRDRs proteins that shared a common domain RdRP which consist of a sequence motif corresponds to the catalytic β’ subunit of DNA-dependent RNA polymerases [61]. The CsRDRs have the genome length varied from 4373 bp to 11526 bp corresponding coding potentiality of 1157 and 1015 amino acids for CsRDR6 and CSRDR3 respectively.

### 3.2. Phylogenetic Analysis of DCL, AGO and RDR proteins in *C. sinensis* and *A. thaliana*

In order to conduct the phylogenetic relationship analysis among the DCL, AGO and RDR proteins of sweet orange and Arabidopsis, the total length amino acid sequences of these plants were used to assess the evolutionary history throughout the neighbor-joining phylogenetic tree (Fig. **(3)**) construction method with considering 1000 bootstrap values. Phylogenetic tree was generated from the full-length aligned protein sequences (**S**upplementary Data **S1**) of the 4 CsDCLs and 4 AtDCLs from *C*. *sinensis* and Arabidopsis (Fig. **(3A)**).The tree represented four subfamilies of newly identified CsDCL proteins i.e. CsDCL1, CsDCL2, CsDCL3 and CsDCL4 corresponding to the DCLs in Arabidopsis on the basis of high similarity of AtDCL1, AtDCL2, AtDCL3 and AtDCL4 respectively with well-supported bootstrap values. The CsDCL proteins showed high sequence conservation with their corresponding counterpart in Arabidopsis. Every DCL subfamily comprised a single CsDCL protein.

**Fig. (3).**
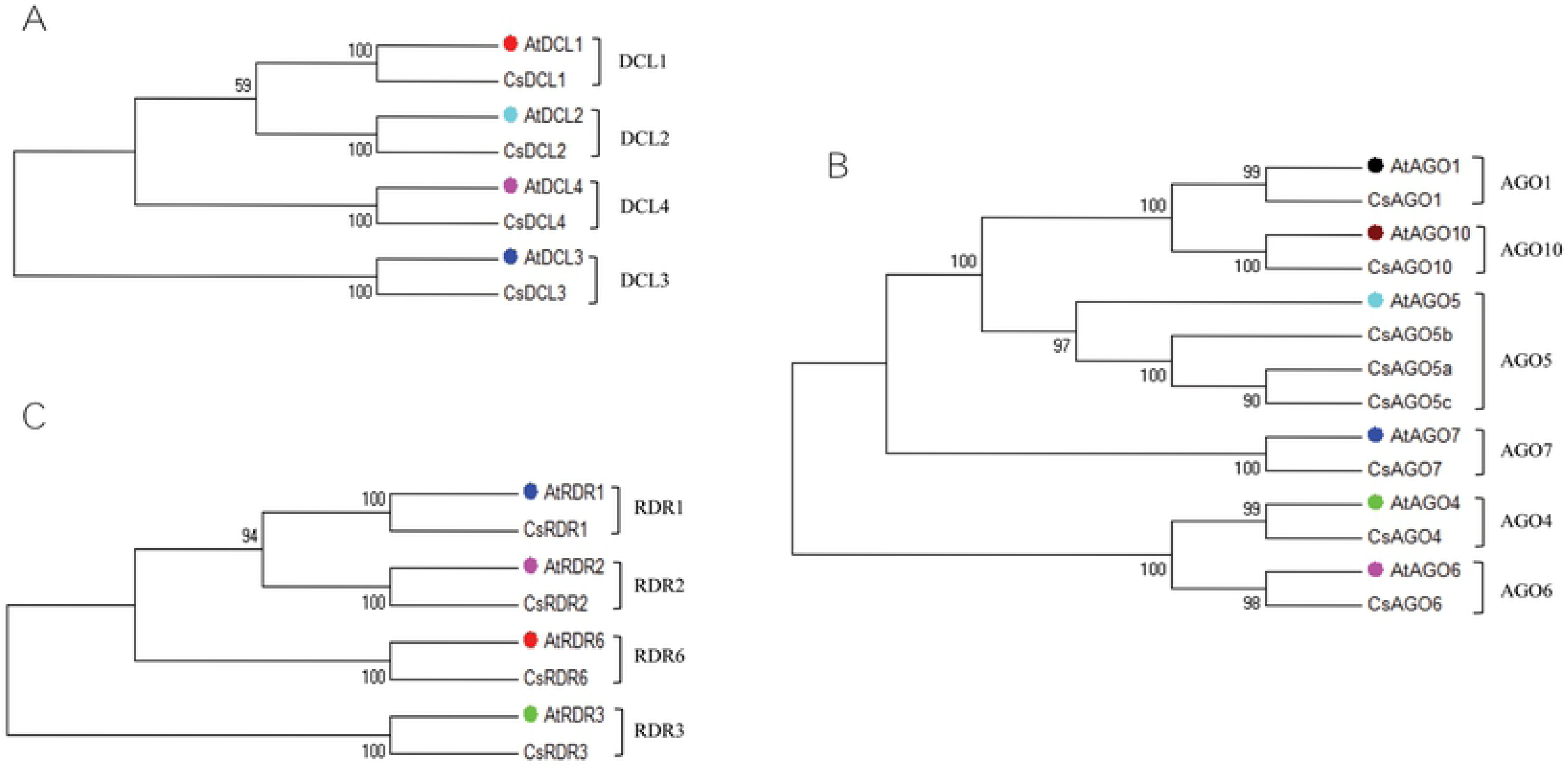
Phylogenetic tree for the (A) Dicer-like (DCL) proteins (B) Argonaute (AGO) proteins and (C) RDR proteins from Citrus and Arabidopsis. All the phylogenetic trees were constructed by neighbour-joining method considering significant bootstrap values. The accession number and the abbreviations of proteins from Arabidopsis are given below while others are tabulated in (Table 1): (A) Four DCL proteins, AtDCL1 (At1g01040), AtDCL2 (At3g03300), AtDCL3 (At3g43920) and AtDCL4 (At5g20320) were used in DCL analysis. (B) AtAGO1 (At1g48410), AtAGO4 (At2g27040), AtAGO5 (At2g27880), AtAGO6 (At2g32940), AtAGO7 (At1g69440) and AtAGO10 (At5g43810) were considered for AGO analysis. (C) Among six AtRDR proteins, the phylogenetic tree exhibited only four major classes with CsRDR proteins. The RDR proteins from Arabidopsis were AtRDR1 (At1g14790), AtRDR2 (At4g11130), AtRDR3 (At2g19910) and AtRDR6 (At3g49500). The three different gene families from Arabidopsis are indicated by coloured in the constructed tree adjacent to the designation.

To construct the phylogenetic tree for CsAGO proteins, the full-length multiple aligned protein sequence of the 8 CsAGOs and 6 AtAGOs were considered (Supplementary Data **S2**). The tree exhibited six subfamilies, AGO1, AGO4, AGO5, AGO6, AGO7 and AGO10 with the higher sequence similarity (Fig. **(3B)**). The AGO1 subfamily consists only a single Citrus protein named CsAGO1 with the Arabidopsis AGO protein AtAGO1. Among the other AGO subfamilies, each showed a group containing a single Citrus AGO protein with a single Arabidopsis AGO protein, except AGO5 cluster.

The AGO5 subfamily included three citrus proteins with a single Arabidopsis protein AtAGO5, which were named CsAGO5a, CsAGO5b and CsAGO5c on the basis of higher sequence similarity to AtAGO5. The AGO1, AGO4, AGO6, AGO7 and AGO10; these groups exhibited each separated cluster with a single AGO protein from Citrus and a single protein from Arabidopsis. The clustered citrus AGO proteins were designated as CsAGO1, CsAGO4, CsAGO6, CsAGO7 and CsAGO10 respectively based on high sequence similarity to their corresponding pair AGO protein from Arabidopsis.

Four main classes of RDR genes in Citrus were revealed by the phylogenetic analysis of the full length aligned polypeptide sequences (Supplementary Data **S3**) of RDR proteins of Citrus and Arabidopsis shown in (Fig. **(3C)**). The CsRDR proteins were designated as CsRDR1, CsRDR2, CsRDR3 and CsRDR6 corresponding to the RDR proteins of Arabidopsis AtRDR1, AtRDR2, AtRDR3 and AtRDR6 respectively for the increased sequence similarity. The predicted CsRDR proteins were clustered according to their high sequence conservation with their reflection part in Arabidopsis RDR proteins.

### 3.3. Conserved Domain and motifs analysis of predicted proteins

Preliminary domain analysis of the predicted CsDCLs, CsAGOs and CsRDRs were conducted for searching the conserved domains by Pfam databases, NCBI-CDD databases and Simple Modular Architecture Research Tool (SMART) analysis, results are tabulated in (Table **(3)**).

**Table 3:** Domain analysis of the DCLs, AGOs and RDRs proteins of the predicted gene mapping on *C*. *sinensis* with Pfam, SMART and NCBI-CDD.

| CsDCLs |  |  |  |  |  |
| --- | --- | --- | --- | --- | --- |
| Serial Number | Gene Name | Accession Number | Domains |  |  |
|  |  |  | Pfam | SMART | NCBI-CDD |
| CsDCLs |  |  |  |  |  |
| 1 | CsDCL1 | orange1.1g000174m | DEAD, Helicase_C, Dicer_dimer,PAZ, Ribonuclease_3,Ribonuclease_3, dsrm, DND1_DSRM | DEXDc, Helicase_C, Dicer_dimer, PAZ, RIBOc, RIBOc, DSRM, DSRM | PAZ super family, helicase_C, Rnc, RIBOc, Dicer_dimer, DSRM |
| 2 | CsDCL2 | orange1.1g000607m | DEAD, Helicase_C, Dicer_dimer, PAZ, Ribonuclease_3, Ribonuclease_3 | DEXDc, Helicase_C, Dicer_dimer, PAZ, RIBOc, RIBOc | helicase_C, RIBOc , PAZ super family, Dicer_dimer, RIBOc |
| 3 | CsDCL3 | orange1.1g000379m | ResIII,Helicase_C, Dicer_dimer, PAZ, Ribonuclease_3 | DEXDc, Helicase_C, Dicer_dimer, Low complexity PAZ, RIBOc | helicase_C, PAZ super family ,RIBOc, Dicer_dimer, Rnc super family |
| 4 | CsDCL4 | orange1.1g000380m | DEAD, Helicase_C, Dicer_dimer, PAZ, Ribonuclease_3, Ribonuclease_3, DND1_DSRM | DEXDc, Helicase_C, Dicer_dimer, PAZ, RIBOc, RIBOc, DSRM, DSRM | helicase_C, RIBOc, Dicer_dimer, RIBOc, PAZ super family, DSRM |
| CsAGOs |  |  |  |  |  |
| 1 | CsAGO1 | orange1.1g001466m | Gly-rich_Ago1, ArgoN, ArgoL1, PAZ, ArgoL2, ArgoMid, Piwi | Gly-rich_Ago1, ArgoN, DUF1785, ArgoL1, PAZ, ArgoL2, ArgoMid, Piwi | Piwi-like super family, Gly-rich_Ago1 |
| 2 | CsAGO4 | orange1.1g002449m | ArgoN, ArgoL1, PAZ, ArgoL2, ArgoMid, Piwi | ArgoN, DUF1785, PAZ, ArgoL2, ArgoMid, Piwi | Piwi-like super family |
| 3 | CsAGO5a | orange1.1g002204m | ArgoN, ArgoL1, PAZ, ArgoL2, ArgoMid, Piwi | ArgoN, DUF1785, PAZ, ArgoL2, ArgoMid, Piwi | Piwi-like super family |
| 4 | CsAGO5b | orange1.1g037086m | PAZ, ArgoL2, ArgoMid, Piwi | PAZ, ArgoL2, ArgoMid, Piwi | Piwi-like super family, PAZ |
| 5 | CsAGO5c | orange1.1g003630m | ArgoN, ArgoL1, PAZ, ArgoL2, ArgoMid, Piwi | ArgoN, DUF1785, PAZ, ArgoMid, Piwi | Piwi-like super family |
| 6 | CsAGO6 | orange1.1g002661m | ArgoN, ArgoL1, PAZ, ArgoL2, Piwi | ArgoN, DUF1785, PAZ, ArgoL2, Piwi | Piwi-like super family |
| 7 | CsAGO7 | orange1.1g001684m | ArgoN, ArgoL1, PAZ, Piwi | ArgoN, DUF1785, PAZ, ArgoMid, Piwi | Piwi_ago-like, PAZ, ArgoL1, ArgoN |
| 8 | CsAGO10 | orange1.1g001954m | ArgoN, ArgoL1, PAZ, ArgoL2, ArgoMid, Piwi | ArgoN, DUF1785, PAZ, ArgoL2, ArgoMid, Piwi | Piwi-like super family |
| *CsRDRs* |  |  |  |  |  |
| 1 | CsRDR1 | orange1.1g002586m | RdRP | RdRP | RdRP |
| 2 | CsRDR2 | orange1.1g001183m | RdRP | RdRP | RdRP, RRM_SF super family |
| 3 | CsRDR3 | orange1.1g001771m | RdRP | RdRP | RdRP |
| 4 | CsRDR6 | orange1.1g041430m | RdRP | RdRP | RdRP |

The CsDCLs proteins showed all the conserved domains through the SMART analysis exhibited some unknown regions and low complexity regions besides the expected domains (Fig. **(4)**). The conserved motifs were screened using MEME with the pre-discussed criterions. The CsDCLs, CsAGOs and CsRDRs proteins were performed all characteristic of DCL, AGO and RDR domain organization was found in Arabidopsis DCLs, AGOs and RDRs proteins respectively when the CsDCL3 has the ResIII domain instead of DEAD domain and CsAGO5b has missed the ArgoN domain having the PAZ and PIWI domain (Fig. **(4)**).

**Fig. (4).**
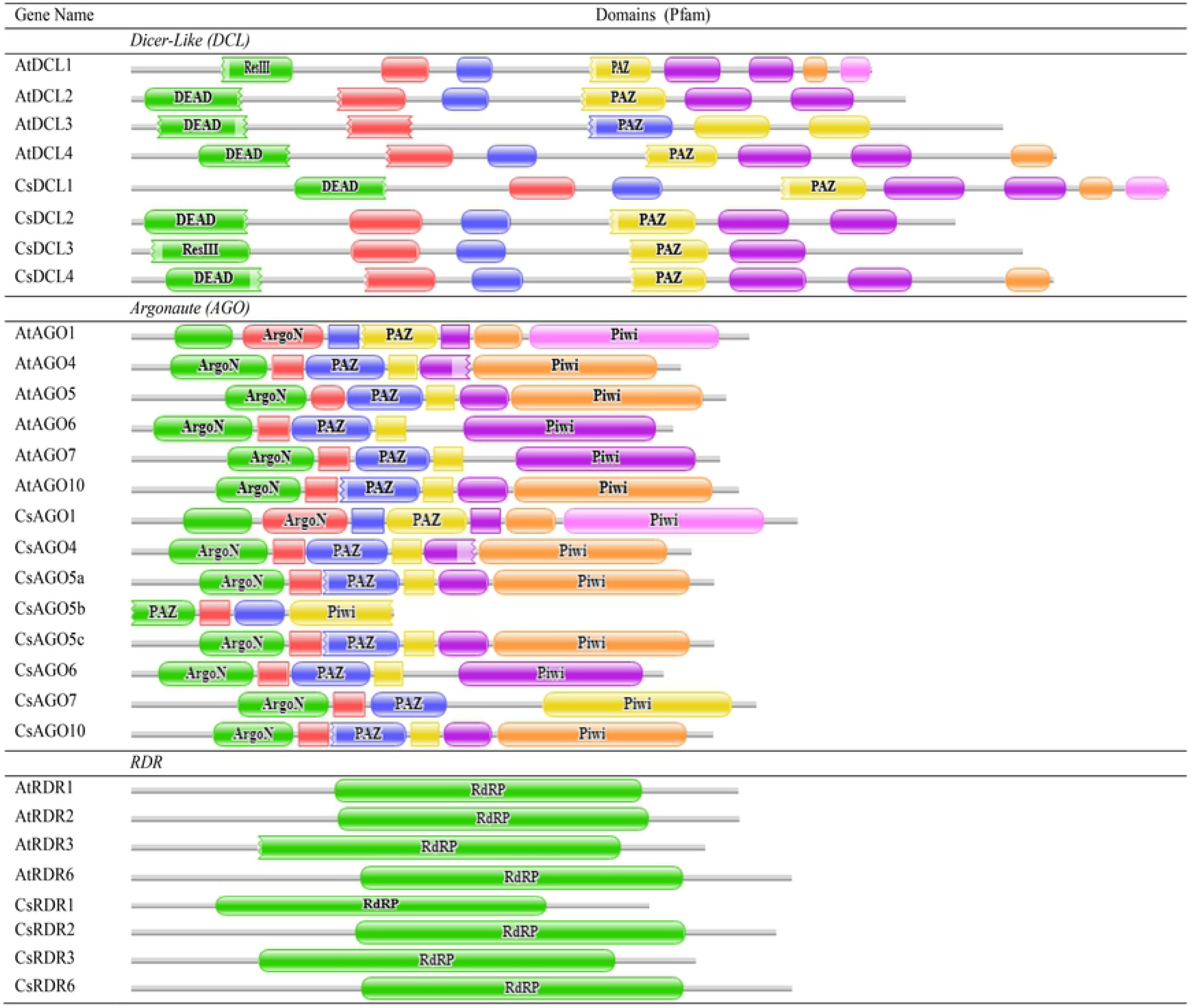
The conserved domains of the predicted proteins were drawn by using Pfam database.

By MEME-suite analysis, the DCLs proteins have 19 (in CsDCL2 and CsDCL3) and 20 (in CsDCL1 and CsDCL4) motifs among the 20 motifs as mentioned before. The predicted motifs are well distributed among the DCL domains for all CsDCLs proteins The MEME analysis of CsAGOs proteins identified 16 common conserved motifs from 20 motifs among all the AGO proteins in *C*. *sinensis* and Arabidopsis, except the CsAGO5b having 9 conserved motifs. The predicted conserved motifs were distributed among the AGO domains in *C*. *sinensis* AGO proteins.

Among the CsAGO proteins one has 16 different motifs (CsAGO7), three proteins (CsAGO4, CsAGO5c and CsAGO6) reflected 17 motifs and others three proteins (CsAGO1, CsAGO5a and CsAGO10) contained 20 conserved motifs (Fig. **(5)**). Although from the analysis, it is observed that the well conservation within the AGO proteins of *C*. *sinensis* and Arabidopsis, there still has had some variability of motif distribution between the different subfamilies of AGO proteins in *C*. *sinensis.* Moreover, this analysis suggested that the conserved predicted motifs might play some vital functional roles in these AGO proteins.

**Fig. (5).**
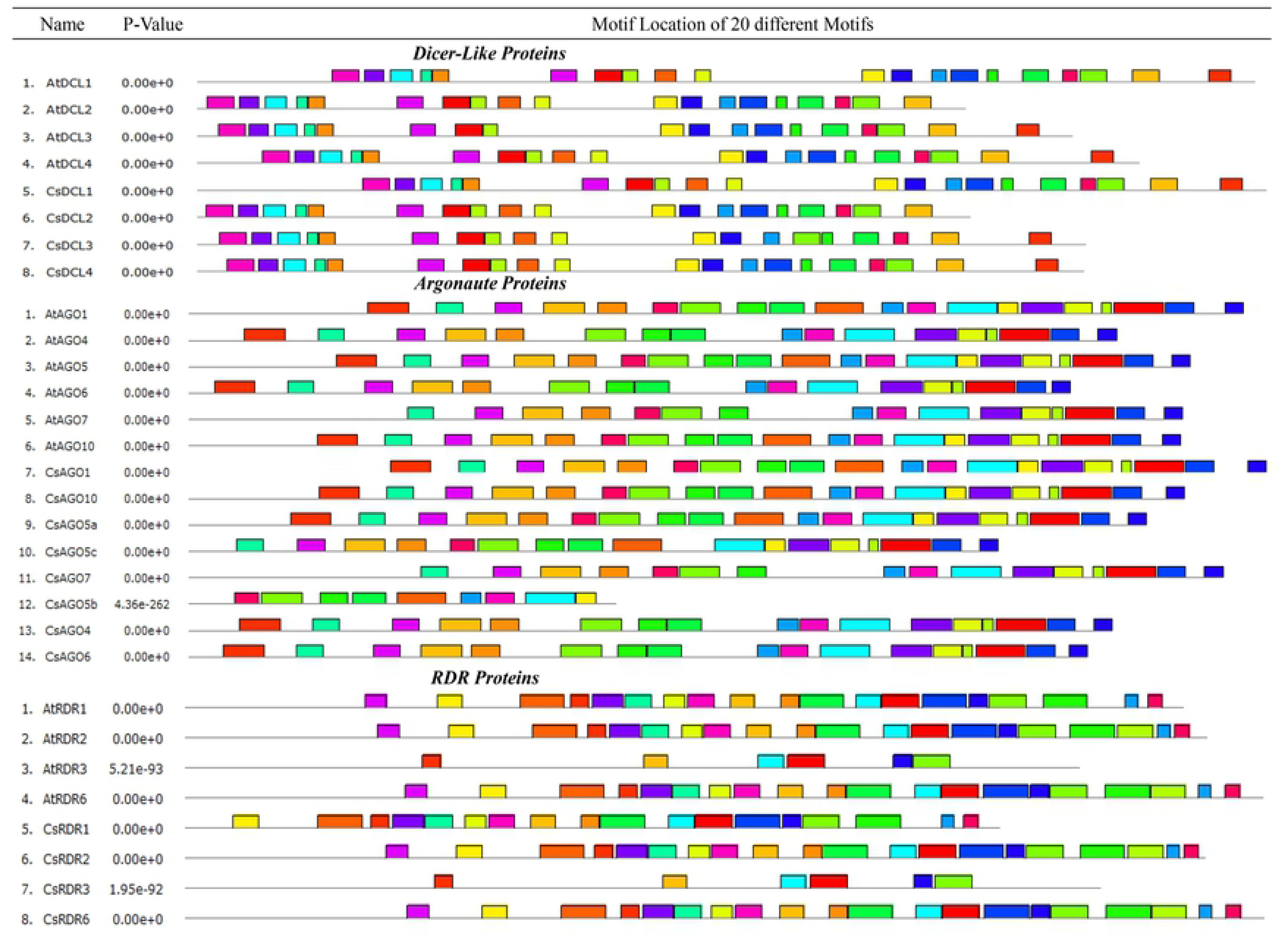
Conserved motifs of the proteins of different genes families are drawn by MEME-suite (maximum 20 motifs are displayed). Different color represented various motifs distributed in the domains of the proteins.

In RDR protein family, the MEME analysis exhibited that the least 6 conserved motifs in CsRDR3 coincided with the AtRDR3. Among other CsRDRs proteins, the CsRDR2 and CsRDR6 lighted 20 out of 20 conserved motifs which are well distributed on the RdRP domain and the CsRDR1 had 18 out of 20 conserved motifs (Fig. **(5)**). Although the predicted motifs were well conserved in the major part of the RDR domain, the motif schemes of different RDR subfamilies did not follow the same distributional pattern. The RDR proteins also reflected some additional motifs besides the RdRP domain having the unknown functional role. However, the MEME-suite analysis reflexed that the CsDCLs, CsAGOs and CsRDRs proteins are enriched with well conservation and distribution of the motifs throughout the subfamilies. This analysis suggested that the predicted motifs might play a different vital role in the functional importance of these genes in *C*. *sinensis* in different life stages.

### 3.4. Gene Structure and Genomic location of CsDCLs, CsAGOs and CsRDRs

To observe the gene structure of the predicted CsDCLs, CsAGOs and CsRDRs gene, their exon-intron configuration was explored by using GSDS with the respective genes family of Arabidopsis. The exon-intron configuration of the predicted genes represented higher conservation as expected for that of DCLs, AGOs and RDRs genes in the model plant Arabidopsis (Fig. **(6)**). The gene structure of CsDCLs exhibited having the number of intron 18-25 (Table **(1))**, correspondence according to higher similarity with AtDCLs.

**Fig. (6).**
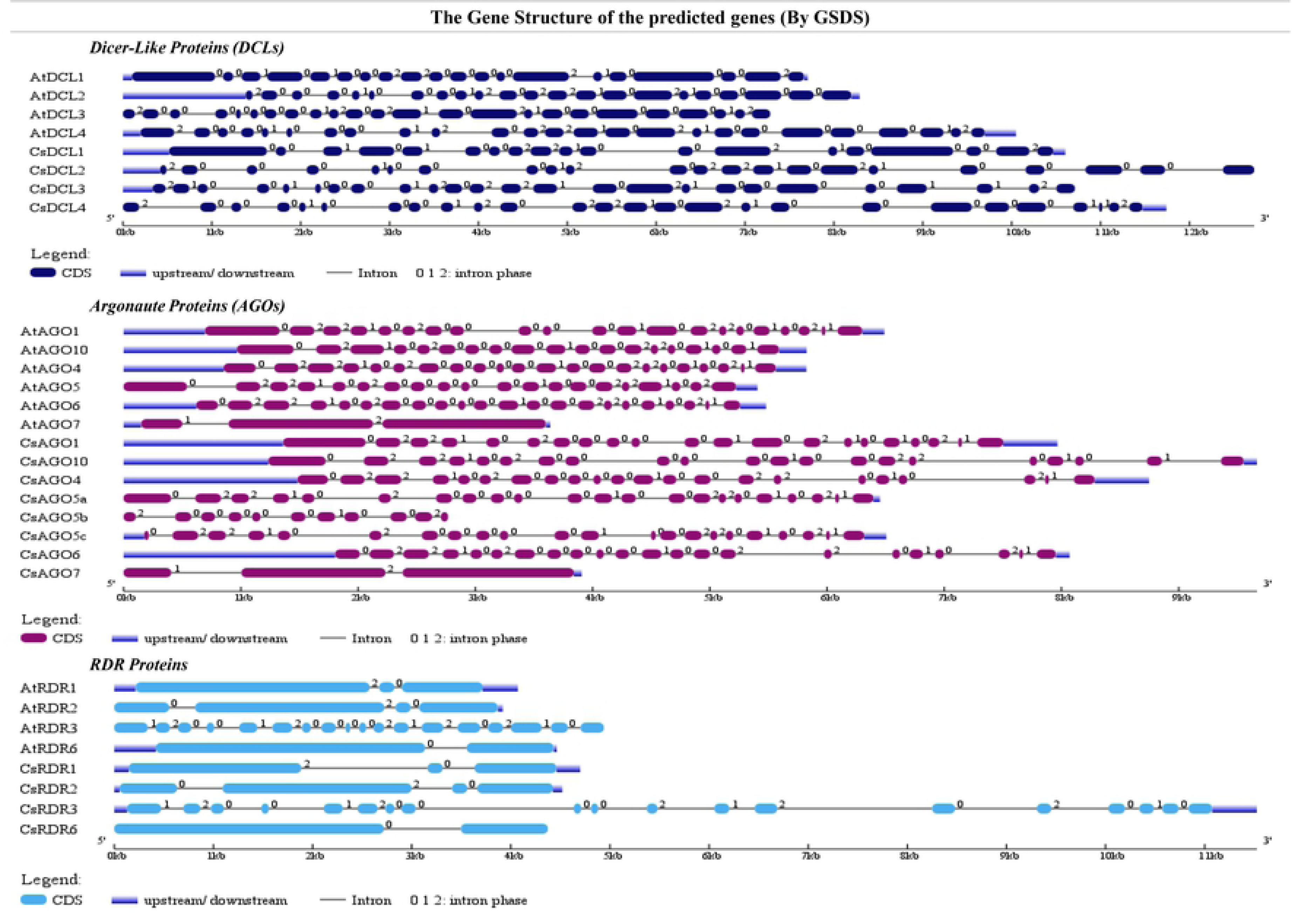
Gene structure of the predicted CsDCLs, CsAGOs and CsRDRs proteins in *C. sinensis* with Arabidopsis using Gene Structure Display Server (GSDS 2.0, http://gsds.cbi.pku.edu.cn/index.php) (Bo Hu et al. 2015).

On the other hand, out of eight CsAGOs, six genes displayed 20 or 21 introns in the gene structure except the CsAGO5b and CsAGO7 having the number of introns 10 and 2 respectively (Fig. **(6)**). This structure indicated that CsAGOs genes are highly similar to the AtAGOs. The CsRDRs showed up with the equal number of introns with their paralogs from Arabidopsis, except the CsRDR3 having a number of intron 18 which is just one short than introns in AtRDR3.

The genomic location of the predicted RNAi pathway related genes in *C*. *sinensis* was conducted by observing the position of the genes in different scaffold location. The predicted CsDCLs, CsAGOs and CsRDRs genes were distributed among the 15 different scaffolds through the entire genome (Supplementary Table **S1**). All proteins have a unique scaffold position while the two CsAGO genes (CsAGO5a and CsAGO5b) were placed in the scafold000595 in the different location. Most of the genes are situated from scafold00001 to scafold00674 where the CsAGO5c is in the scafold03700.

### 3.5. Gene Ontology Enrichment Analyses

In order to better understand the biological roles of the predicted RNAi genes and characterize them, GO enrichment analysis was performed (Fig. **(7)**, Supplementary Data **S4**). From the analysis result it was observed that 12 genes involved in post-transcriptional gene silencing (PTGS) pathway (GO:0016441; p-value:3.60e-27), 10 related to RNA interference (GO:0016246; p-value: 8.50e-25) and 12 genes are associated with gene silencing (GO:0016458; p-value:7.40e-24). The RNAi is closely related to the phenomenon named post-transcriptional gene silencing (PTGS) in plants [62].

**Fig. (7).**
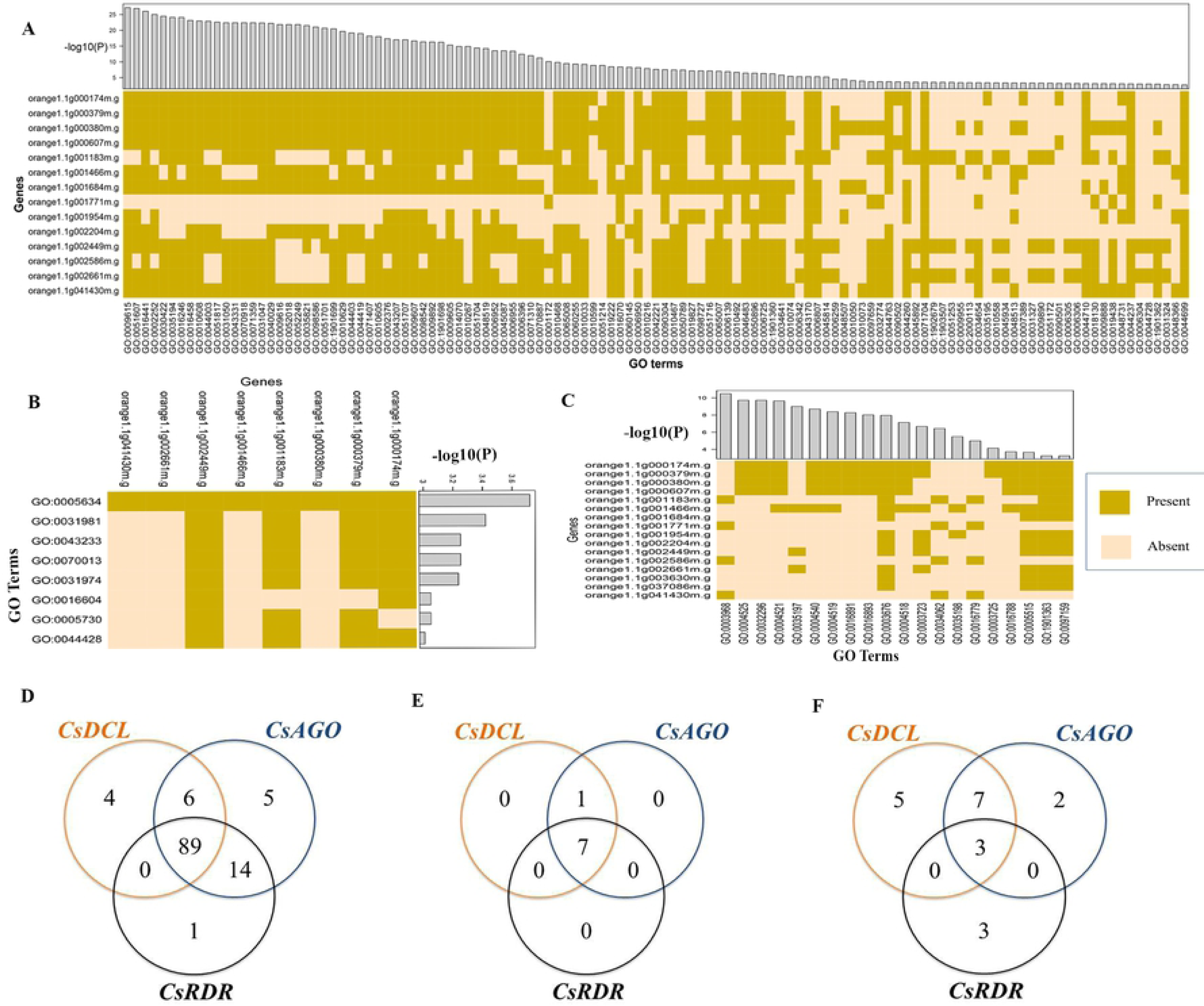
The heatmap for the predicted GO terms corresponding to the RNAi genes are presented for (A) biological process (B) cellular components (C) molecular function whether the genes are related (Present) or not (Absent). The p-value corresponds to the GO terms are showed in histogram adjacent to the heatmap, using -log10 (p-value). The Ven diagrams are drawn to observe the shared GO terms by three gene families considering the (D) biological process (E) cellular components (F) molecular functions.

The GO enrichment analysis showed that five predicted RNAi genes (among 16) are related to the endonuclease activity (GO: 0004519; p-value= 4.20e-09) which (Supplementary Data S4) indicates a positive linkage with the RNA-induced silencing complex (RISC-mediated) mediated cleavage activities into the cell. This multimeric protein complex (i.e. RISC) guides for protein degradation. Among the RNAi proteins, Argonautes work for the cleavage called endonucleolytic activities, which result the final PTGS for specific mRNA substrate [63]. By the GO analysis, there are 13 genes related to nucleic acid binding (GO: 0003676; p-value=1.10e-08), 7 genes to RNA binding (GO: 0003723; p-value=2.10e-07) and 12 genes related to protein binding (GO: 0005515; p-value= 0.00022) activities (Supplementary Data S4) which indicate the RNAi protein’s participation to the RISC as well as interference processes. The predicted CsAGO proteins contain the special domains called PAZ and PIWI domain that play the key role in making a complex with RNA or DNA. The PAZ domain has a nucleic acid-binding fold that promotes the domain to bind to the specific position of the nucleic acids [64, 65]. The GO enrichment analysis also showed the attachment of the predicted genes to the numerous biological processes. Significantly, most of the reported genes are engaged with the regulation of biological process (GO:0050789; p-value=3.90e-08), negative regulation of gene expression (GO:0010629; p-value:2.30e-20) and dsRNA fragmentation (GO:0031050; p-value:4.10e-23) (Supplementary Data **S4**).

The *C. sinensis* RNAi genes are also involved in virus response (GO:0009615; p-value:6.70e-28), immune response (GO:0006955; p-value:4.10e-14) as these were reported for AtDCL and AtRDR[11, 21, 33, 39–42]. These GO enrichment analysis for biological processes (Supplementary Fig. **S1A**), molecular functions (Supplementary Fig. **S1B**) and cellular component (Supplementary Fig. **S1C**) undoubtedly indicated that the predicted genes are deeply interrelated with the RNAi pathway in *C. sinensis.* In addition, the predicted genes act with the hydrolase activity, acting on ester bond, predicted from the GO analysis (Supplementary Data **S4**).

The Ven Diagram of the GO terms for three clusters of the RNAi genes was drawn (Fig. **(7)**). It was observed that the CsDCL, CsAGO and CsRDR genes shared significant number of GO pathways in common. In biological process, there are 89 GO enrichment pathways (Fig. **(7)**) were shared by the reported proteins, which indicate the involvement of the RNAi gene members in numerous biological processes together. Also in molecular function and cellular component, the predicted genes exhibited a group of mutual GO pathways. So this GO analysis provides a significant indication of the predicted RNAi genes in this study.

### 3.6. Sub-cellular Localization of the Reported Genes and Proteins

The subcellular localization studies of the predicted proteins were observed to uncover their cellular appearance. By the sub-cellular localization annotation, it has been shown that all the reported proteins identified in this study appear into the cytosol (Fig. **(8)**). As PTGS occurs into the cytoplasmic region [66], this result implies that the reported RNAi proteins may directly involve to the PTGS process. On the other hand, four CsAGO and one CsRDR proteins exhibited their appearance into the nucleus when as no CsDCLs are located there. These bring a significant importance whether the CsDCLs are not found in nucleus. Further expression pattern analysis will provide deeper information about the CsDCLs. Some of the identified RNAi proteins are also distributed into the cell membrane, plastid and mitochondria (Fig. **(8)**).

**Fig.(8).**
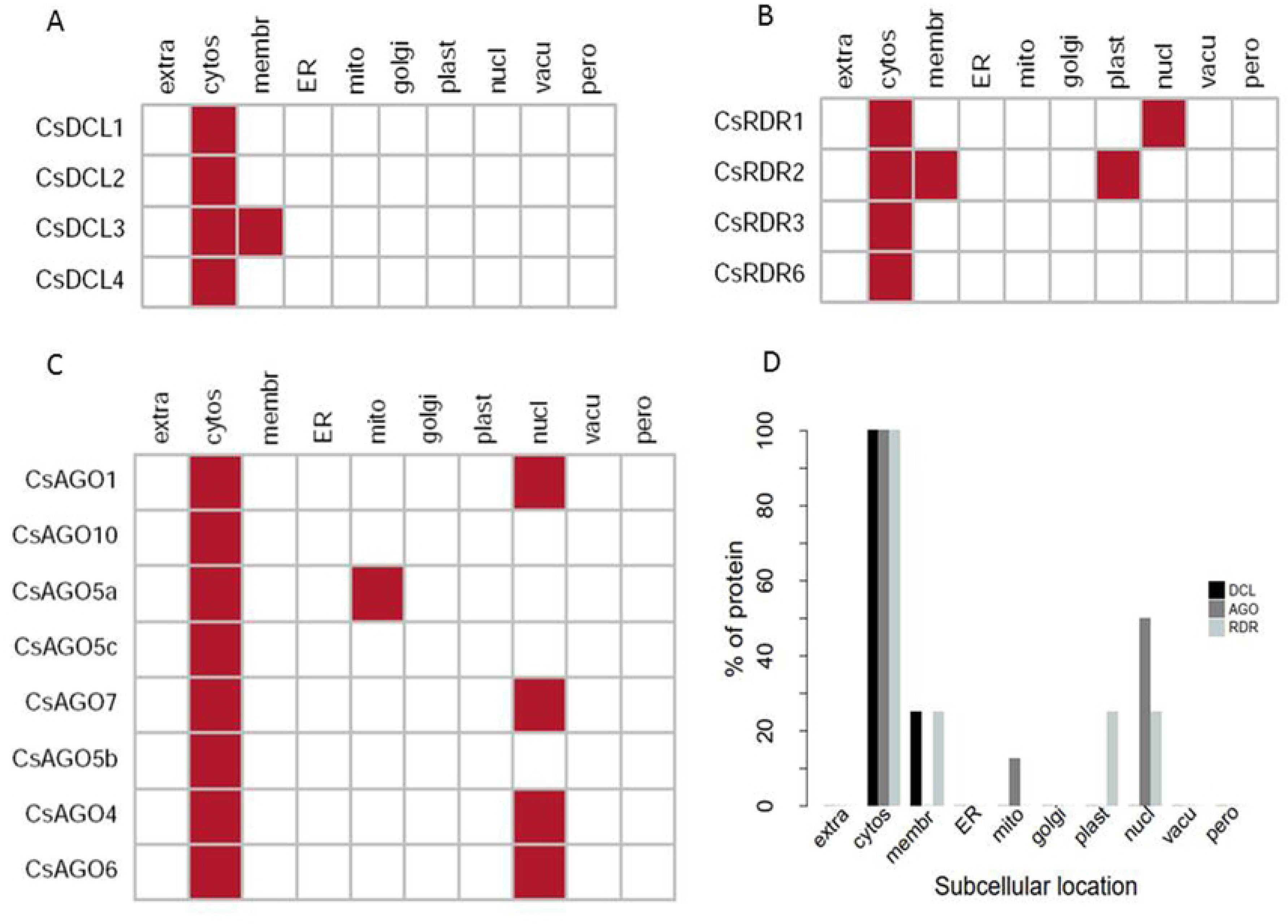
Sub-cellular localization analysis for (A) CsDCL (B) CsRDR and (C) CsAGO proteins. (D) The percentage of protein appears in different cellular components. Here cytosol (cytos), endoplasmic reticulum (ER), extracellular (extra), golgi apparatus (golgi), membrane (membr), mitochondria (mito), nuclear (nucl), peroxisome (pero), plastid (plast) and vacuole (vacu). Over all report is tabulated in (Supplementary Table **S2**).

Previous studies reported that the RNAi genes are not only highly related with PTGS but also with transcriptional gene silencing (TGS) [66]. In protein transcriptional process, RNA polymerase type II complexes are directly involved [67]. For PTGS, the RNAi proteins have greater participation in RNA-induced silencing complex (RISC-mediated) mediated cleavage activities by the help of DCL, AGO and RDR proteins with other molecules [63]. The PTGS happens into the cytoplasmic region for targeted mRNA protein degradation [67].

### 3.7. Regulatory Relationship between TFs and RNAi genes

TFs are considered as the switch of gene expression in all living organism. Plant TFs play significant role in growth, development and stress response activities [68]. Thus, identifying TFs regulating RNAi genes could help to improve our understanding of gene silencing process in *C. sinensis*. In this analysis a total of 235 TFs were identified those regulate the predicted RNAi genes (Supplementary Data **S5**). The identified TFs were distributed into 27 groups based on the TF families. The TFs MYB, Dof, ERF, NAC, MIKC_MADS, WRKY and bZIP families might play significant role in regulating RNAi genes. Particularly those of ERF, NAC, WRKY and bZIP families, which are the top four families those contained 29, 20, 20 and 10 TFs and accounted 57.66% of the total identified TFs (Supplementary Table **S3**). This finding indicated that those TFs could be important in regulating RNAi genes.

To better understand the regulatory relationship between TFs and RNAi genes, a regulatory network was constructed (Fig. **(9A)**). From the resultant network it is observed that different groups of TFs exhibited distinct structure. For example, TF belongs to ERF family mainly bind to the gene CsAGO5a (Fig. **(9B)**, Supplementary Fig. **S2**). However some RNAi genes such as CsAGO5c, CsDCL4 and others are regulated by ERF family; all of them are also linked to CsAGO5a (Fig. **(10A)**). Very similar results were also observed for the hub TFs NAC, WRKY and bZIP (Fig. **(10B-D)**).

**Fig.(9).**
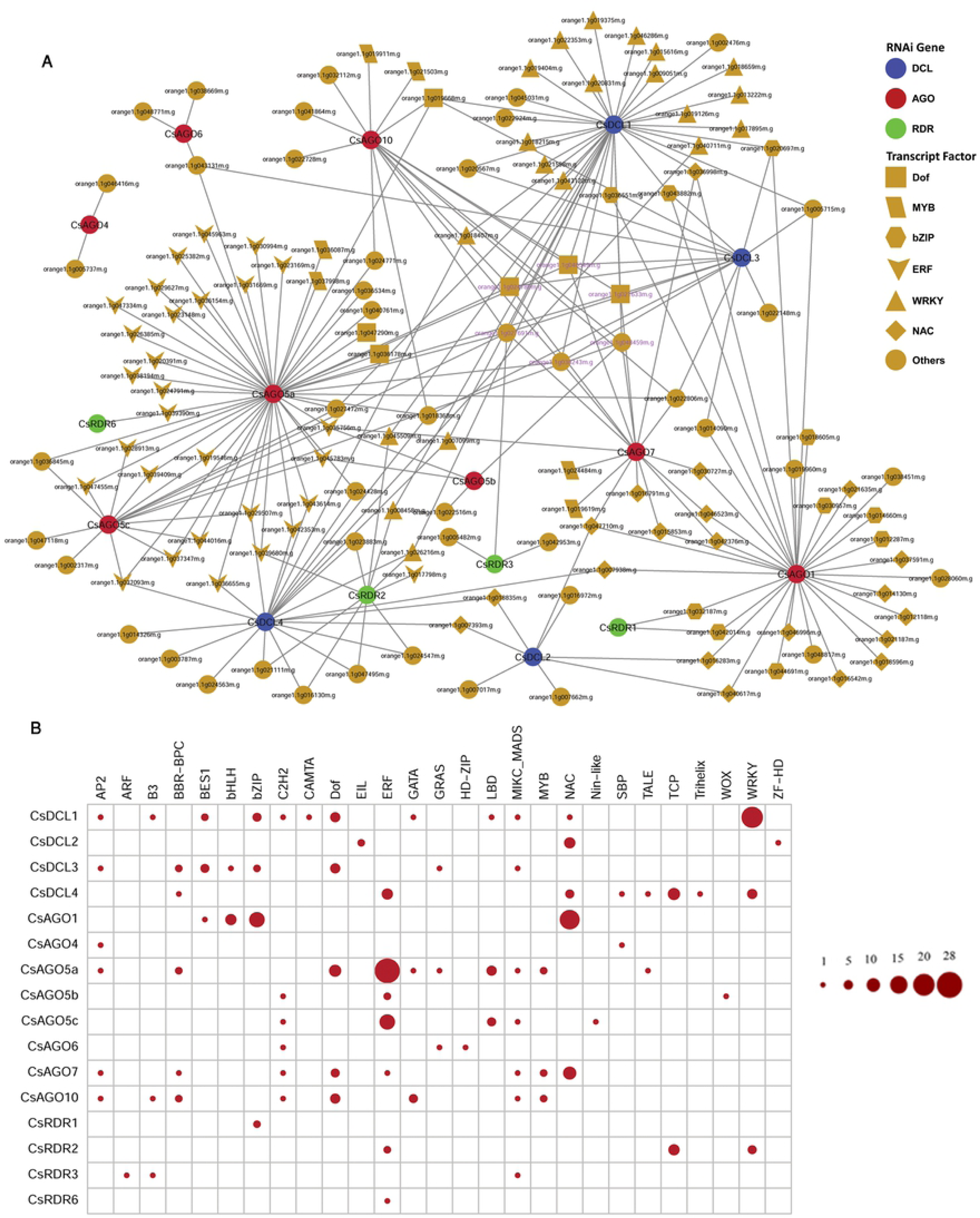
**(A)** The regulatory network among the TFs and the predicted RNAi genes. The nodes of the network were colored based on RNAi genes and TFs. DCL, AGO and RDR genes were represented by blue, red and green node color respectively and the TFs were represented by yellow node color. Different node symbols were used for different families of TFs. Magenta node level were used for the hub TFs. **(B)**The map representing the associated number of TFs with the CsRNAi genes.

**Fig.(10).**
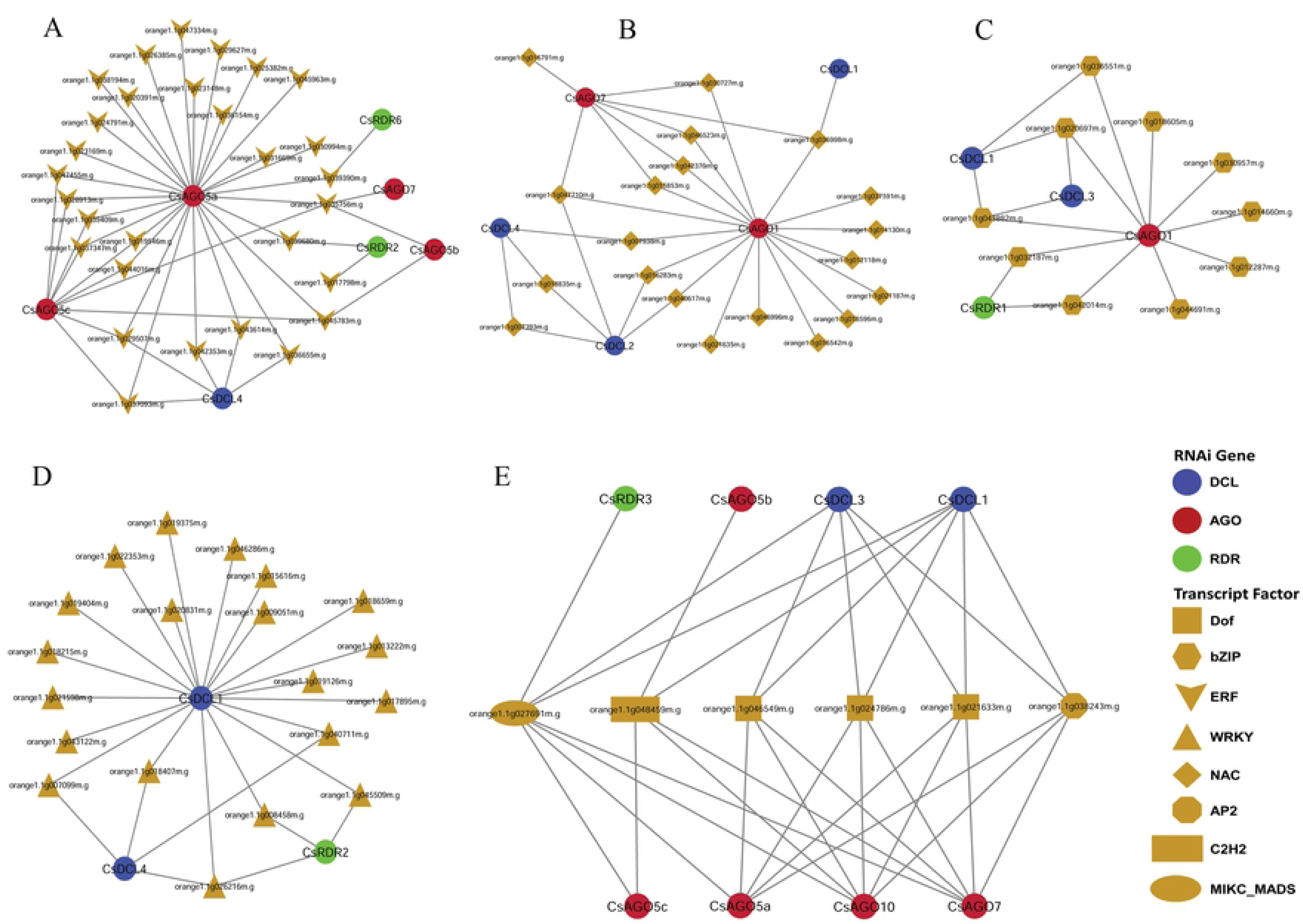
RNAi gene mediated sub-network for (A) ERF, (B) NAC, (C) bZIP, and (D) WRKY TF family. (E) Sub-network among the hub TFs those regulate more than five RNAi genes.

Moreover, six hub TFs were identified on the basis of node degree which have more than five interacting partners with the RNAi genes. Among them 3 belongs to dof family and other three are in MIKC_MSDS, C2H2 and bZIP (Fig. **(10E)**). The Dof TFs family is directly involved with the DNA binding activities by the N- and C-terminal region and causes the regulation of gene activation or repression of the target genes which is the main theme of RNAi. The Dof TFs family also works for the biosynthesis of flavonoids and glucosinolates, stress tolerance, seed germination and controlling the photoperiodic flowering [69–72]. The TF orange1.1g027691m (MIKC_MADS) regulates heights seven RNAi genes and the rest regulate five RNAi genes in the network (Fig. **(10E)**). The regulatory network clearly exposes that these predicted genes of RNAi process in *C. sinensis* will exhibit a dramatically expression pattern that can be retrieved by deeper investigation of these genes in future.

Besides, all of the hubs TFs are connected to eight RNAi genes. Of RNAi genes corresponding to hub TFs, five are AGO two are DCL and only one is RDR. Three RNAi genes (2 AGO: CsAGO10, CsAGO7; 1 DCL: CsDCL1) were predicted to be regulated by the entire six hub TFs.

#### *Cis*-acting regulatory element Analysis

The *cis*-acting regulatory element analyses were conducted to find out the functional diversity of the motifs related to the promoter region of the proposed RNAi genes into *C. sinenesis.* The PlantCARE database provided the information about the motifs and their functionality with the genes. The analysis revealed that most of the motifs were light responsive (LR) (Fig. 12), widely presents in the entire RNAi gene’s promoter. Supporting the EST analysis, the light responsiveness is associated with the photosynthesis which occurs in leaf. Among the light responsive motifs the ATCT-motif, ATC-motif, Box-4, AE-box, G-box, I-box, GAT-motif, GT1-motif were shared by the most of the RNAi genes in *C. sinensis* (Fig. 12). The TC-rich repeats (cis-acting element involved in defense and stress responsiveness), MBS (MYB binding site involved in drought-inducibility) and LTR elements (*cis*-acting element involved in low-temperature responsiveness) were commonly found as stress responsive motif among the CsDCL/AOG/RDR genes in *C. sinensis*. It is known that the plant hormones are essential for plant growth and development.

The significant plant hormone responsive (HR) *cis*-acting element were identified in this analysis. The ABRE (cis-acting element involved in the abscisic acid responsiveness), AuxRR-core (cis-acting regulatory element involved in auxin responsiveness), GC-motif (enhancer-like element involved in anoxic specific inducibility), GARE-motif (gibberellin-responsive element), O2-site (cis-acting regulatory element involved in zein metabolism regulation), P-box (gibberellin-responsive element), TATC-box (cis-acting element involved in gibberellin-responsiveness), TCA-element (cis-acting element involved in salicylic acid responsiveness) and TGA-element (auxin-responsive element) were the hormone responsive *cis*-element shared by the CsDCLs, CsAGOs and CsRDRs as phytohormones responsiveness element (Fig. 12). There are some others significant elements were identified and represented as others activities. The AT-rich element (binding site of AT-rich DNA binding protein (ATBP-1)), AT-rich sequence (element for maximal elicitor-mediated activation), CAAT-box (common cis-acting element in promoter and enhancer regions), CAT-box (cis-acting regulatory element related to meristem expression), CCAAT-box (MYBHv1 binding site), GCN4_motif (cis-regulatory element involved in endosperm expression), TATA-box (core promoter element around −30 of transcription start), circadian (cis-acting regulatory element involved in circadian control), silencer (GT-1 factor binding site) and TGACG-motif (cis-acting regulatory element involved in the MeJA-responsiveness) were recognized as others cis-acting regulatory element shared by RNAi genes in *C. sinensis* (Fig 12). Some unknown cis-regulatory elements were also detected along with the reported cis-elements (Supplementary Table S6). In general the cis-regulatory elements carried out significant evidences about the proposed RNAi genes that will be helpful for further investigation about their role in plant disease response, growth and development.

#### *In silico* Expressed Sequence Tag (EST) Analysis

For the important and valuable information about the gene expression, the expressed sequence tag (EST) data analysis can bring a significant piece. This analysis helps to validate the genes activities and to investigate the activities in various stress conditions. The EST mining results from the PlantGDB database indicated that the RNAi genes of *C. sinensis* expressed in multiple important tissue and organs. However, the expression evidence of the RNAi genes in several species has been reported which showed that the RNAi genes have expression in leaf, root, flower, seeds and in others tissue [13, 23–30, 73]. The *in silico* expressed sequence tag (EST) analysis of the proposed *C. sinensis* RNAi genes showed that all the genes have their expression at least in one organ or tissue. In general, most of the genes were expressed in leaf and fruit which indicate that these genes have a direct involvement with the photosynthesis and fruit developmental stages in *C. sinensis* (Fig. 11).

**Fig.(11).**
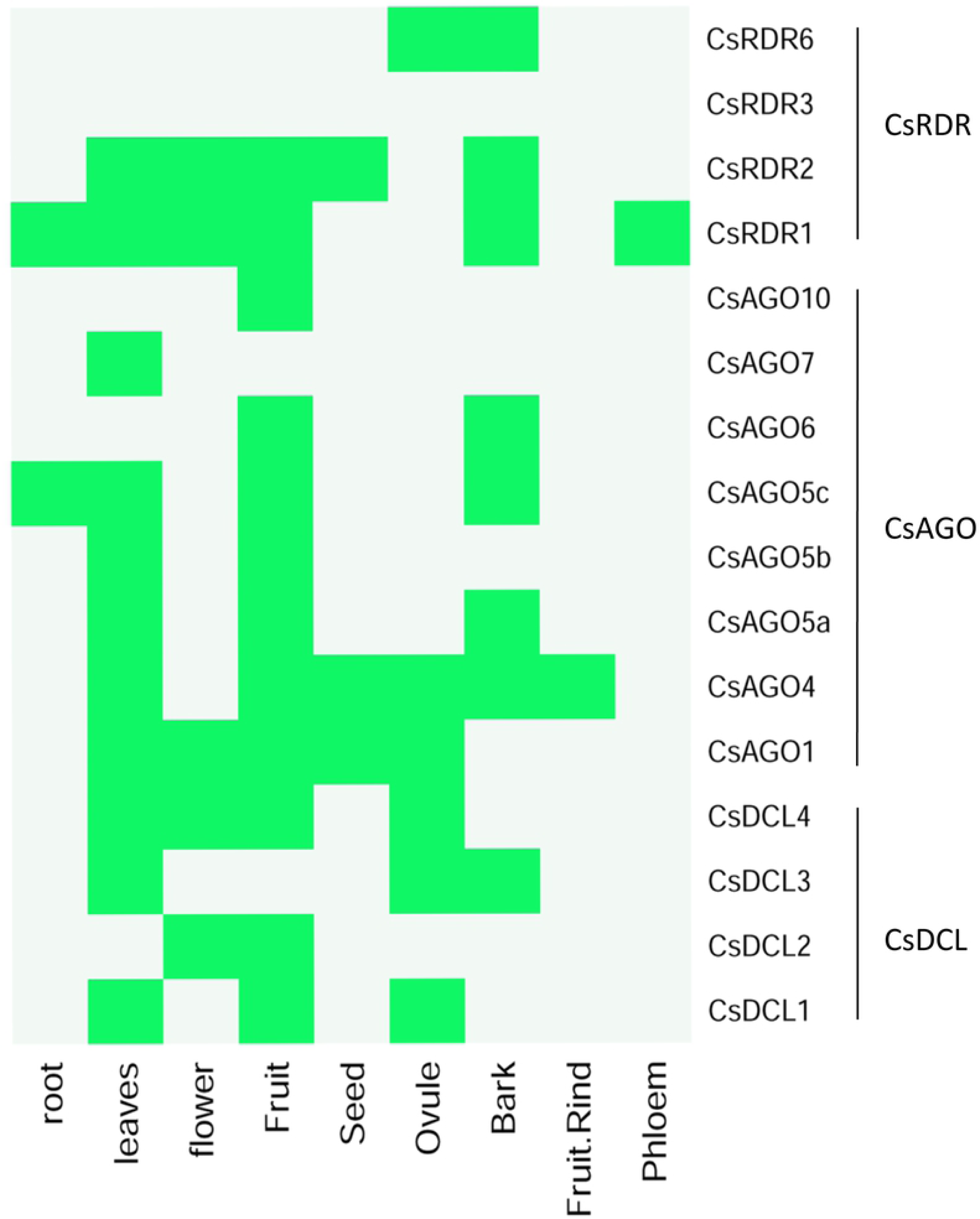
The *in silico* expressed sequence tag (EST) analysis of the identified RNAi genes in C. sinensis plant. The green color represents the existence of expression and off color stands for absent of expression.

**Fig.(12).**
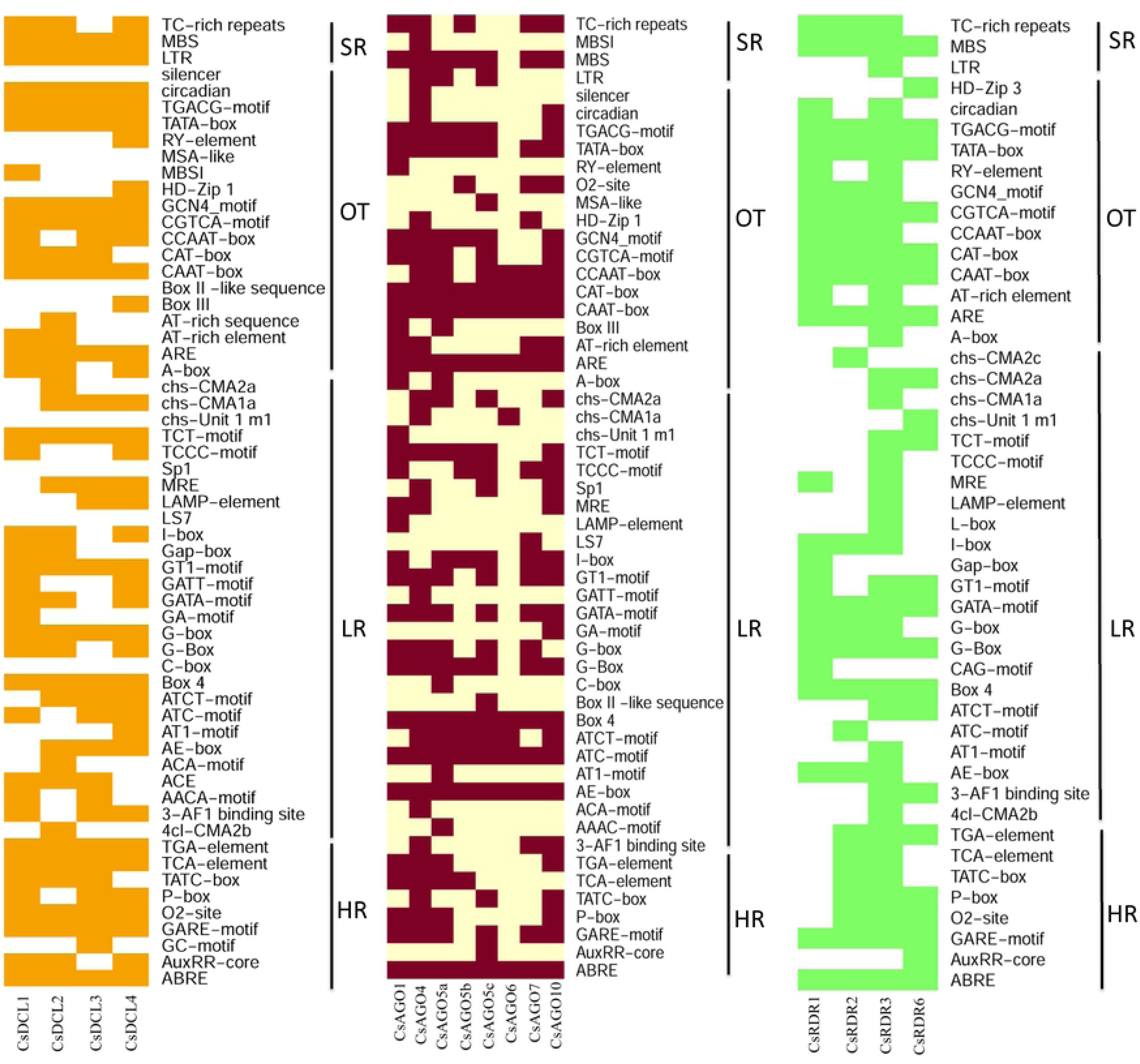
The *cis*-regulatory element in the promoter region of the identified *C. sinenis* DCLs, AGOs and RDRs genes respectively. The deep color represents the presence of that element with the corresponding genes.

The expression of CsDCLs were identified in leafs (CsDCL1/3/4), flowers (CsDCL2/4), fruit (CsDCL1/2/4), ovule (CsDCL1/3/4) and bark (CsDCL3), where no expression was found in root. The entire CsAGOs exhibited diverse expression pattern all over the plant through root, leafs, flowers, ovule, fruit, fruit rind and seed. Among the CsAGO genes, CsAGO1 and CsAGO4 were detected in maximum organs in *C. sinensis.* The only CsAGO1/4 provided the expression evidence into the seeds. Similarly the others RNAi genes, the CsRDRs also highly expressed into the leaf, flower and fruit when the CsRDR6 showed expression in ovule and bark. Interestingly, no expression evidence was found for the CsRDR3 in this *in silico* EST analysis (Fig. 11). According to the EST analysis, the proposed RNAi genes have vast contribution in ovule fertilization, fruit development process, plant photosynthesis which can be dugout by experiment expression validation.

## 4. Conclusion

The orange is considered as the second highest produced fruits all over the world. The *C*. s*inensis* plant is the major source of sweet orange which is one of the most favourite and nutritious fruits. Identification, characterization and diversity analyses of the RNAi gene families were essential, since these families play the vital role for silencing of other gene families of any plant. In this study, a number of bioinformatics approaches were used to identify, characterize and analyse the diversification, properties and biological functionality of RNAi genes in *C*. *sinensis*. By this ways, we identified 4 CsDCL, 8 CsAGO and 4 CsRDR genes as the RNAi gene families in sweet orange. This study also provided additional genomic and physicochemical information of the predicted genes and corresponding proteins. With the phylogenetic analysis, the subgroups of the three gene families were exhibited, the domains and motifs configuration and the gene structures revealed the maximum homogeneity with the respective gene family of Arabidopsis. Moreover, the GO enrichment and subcellular localization analysis provided the final confirmation about the reported genes and protein that, these are the key factor of RNAi process in *C*. *sinensis*. In this analysis, we explored regulatory relationship network between TFs and proposed RNAi genes. Potential targets of TFs were identified those involved in plant growth and development as well as controlling the gene expression or suppression related to RNAi process. The *cis*-acting regulatory elements and expressed sequence tag (EST) analysis indicates that our reported RNAi genes have diverse involvement into the orange plant growth, flowering, and development processes. Thus the reported genes in this study may exhibit significant expression pattern under different stress conditions in various developmental stages of sweet orange like the other plants that were investigated by other researchers. Therefore, the findings of this study may provide a basis for further research on RNAi pathway genes in *C*. *sinensis* to enrich the ultimate sweet orange plant development and fruit production all dover the world.

## Supplementary Files

Supplementary datasets, tables and figures are available at www.bbcba.org/softwares/CsRNAi.zip.

## Abbreviations

AGO: Argonaute
BLAST: Basic Local Alignment Search Tool
bp: base pair
DCL: Dicer-Like
GO: Gene Ontology
GSDS: Gene Structure Display Server
HMM: Hidden Markov Model
MEME: Multiple Em for Motif Elicitation
miRNA: microRNA
PTGS: post-transcriptional gene silencing
RDR: RNA, dependent RNA polymerase
RNAi: RNA interference
RISC: RNA-induced silencing complex
sRNA: small RNA
siRNA: short interfering RNA
TF: transcription factor
TGS: transcriptional gene silencing

## Supportive/supplementary legends

**Supplementary Table S1:** Gene location in different scaffold.

**Supplementary Table S2:** Sub-cellular Localization of the predicted proteins

**Supplementary Table S3:** Distribution of TF families those regulating RNAi genes.

**Supplementary Data S1:** Protein sequences of the predicted DCL genes of *Citrus sinensis*.

**Supplementary Data S2:** Protein sequences of the predicted AGO genes of *Citrus sinensis*.

**Supplementary Data S3:** Protein sequences of the predicted RDR genes of *Citrus sinensis*.

**Supplementary Data S4:** GO enrichment analysis result for predicted RNAi genes.

**Supplementary Data S5:** List of transcript factors and their families regulating predicted RNAi genes.

**Supplementary Fig. S1:** GO enrichment analysis of the predicted RNAi genes **(A)** biological process, **(B)** molecular function and **(C)** cellular process. In the directed acyclic graph (DAG) the downstream term corresponds to a subset of the upstream term. The significant (P < 0.05, FDR < 0.05) GO terms are in colored boxes (the degree of color saturation is positively correlated to the enrichment level of the GO term), and non-significant terms are in white boxes.

**Supplementary Fig. S2:** Distribution of TF families corresponding to genes. Rows of the figure represent the predicted RNAi genes and the columns represent the families of the TFs. The number indicates the TF families regulate the RNAi genes.

## Author Contribution

MPM and MNHM designed the research. MPM analyzed the data using different bioinformatics and statistical tools. MAA and MPM carried out the GO enrichment, subcellular localization and regulatory transcription factor analysis. MPM drafted and MNHM finalized the manuscript. HR, ZA, FFA, MMI and MAM attended at the meetings, presentations and discussion regarding the manuscripts. All authors read and approved the manuscript.

